# Spike sorting for large, dense electrode arrays

**DOI:** 10.1101/015198

**Authors:** Cyrille Rossant, Shabnam Kadir, Dan F. M. Goodman, John Schulman, Mariano Belluscio, Gyorgy Buzsaki, Kenneth D. Harris

## Abstract

Developments in microfabrication technology have enabled the production of neural electrode arrays with hundreds of closely-spaced recording sites, and electrodes with thousands of sites are currently under development. These probes will in principle allow the simultaneous recording of very large numbers of neurons. However, use of this technology requires the development of techniques for decoding the spike times of the recorded neurons, from the raw data captured from the probes. There currently exists no practical solution to this problem of “spike sorting” for large, dense electrode arrays. Here, we present a set of novel tools to solve this problem, implemented in a suite of practical, user-friendly, open-source software. We validate these methods on data from rat cortex, demonstrating error rates as low as 5%.

## Introduction

Understanding the computational mechanisms of the brain will require recording from large neuronal populations spread over multiple neuronal structures. One of the most powerful techniques for neuronal population recording is extracellular electrophysiology, whose yield has been greatly boosted by the development of microfabricated electrode arrays ^1-3^. Advances in microfabrication technology have continuously increased the number of recording sites available on neural probes, and the number of recordable neurons is further increased by having closely spaced recording sites ^4^. Indeed, while a single sharp electrode can provide good isolation of one or two neurons, placing as few as four recording sites together in a “tetrode” can reveal the firing patterns of 10-20 simultaneously recorded cells ^5-7^. This increase is possible because each recorded neuron produces extracellular action potential waveforms (“spikes”) with a characteristic spatiotemporal profile across the recording sites ^8-10^. The analytical process of using these waveforms to decipher the firing times of the recorded neurons is known as “spike sorting” ^11, 12^. For tetrodes, analysis of ground truth data obtained with simultaneous intracellular recording, shows that commonly-applied spike sorting methods can have error rates close to an estimated theoretically optimal performance, which varies between recordings but can be as low as 5%^8^.

The most commonly-applied method of spike sorting proceeds in four steps. The first step is spike detection, where action potential waveforms are extracted, typically by high-pass filtering and thresholding. In the second step, each individual action potential waveform is summarized by a compact feature vector, typically using principal component analysis. In the third step, these vectors are divided into groups corresponding to putative neurons using cluster analysis. Although fully automatic spike sorting would be a powerful tool, the output of current algorithms cannot be accepted without human verification ^8^. Thus the fourth step of the spike sorting pipeline is manual inspection of the spike-sorted data, and adjustment of any errors made by automatic algorithms ^13^. A similar situation arises in many fields of data-intensive science: in 3D reconstruction of serial-section electron microscopy data, for example, automatic methods can only be used under the supervision of human operators ^14^. For high-throughput analysis, it is essential that ergonomic software be designed that focuses operator time on only those questions that cannot be solved by the computer alone.

Modern fabrication technology allows for probes with several hundred channels, arranged in groups of up to 32 closely-spaced recording sites for current models ^15, 16^. Spike sorting methods developed for tetrodes do not work for such large, dense electrode arrays. This failure occurs for multiple reasons. First, most tetrode spike detection algorithms are based on the assumption that a spike occurring on any one channel will be seen on the whole shank, which does not hold for shanks of more that ∼8 sites. Second, cluster analysis in high-dimensional spaces is a notoriously challenging statistical problem ^17^, with no simple solution that works in all circumstances. Finally the process of manual verification and adjustment, while manageable with low-count probes, cannot scale to the high-count case without software that guides the operator to only those decisions that cannot be made reliably by a computer. Although many different methods for spike sorting have been proposed (e.g. Refs. ^18-24^); no method has yet solved these pressing problems robustly enough to be widely adopted by the experimental community.

Here we describe a set of novel algorithms and open-source software for the spike sorting of high-channel count electrode data, which are implemented in a suite of three freely available software programs. While the spike sorting problem has attracted considerable theoretical research, our goal here was to produce a practical spike sorting system for high-channel count data, that can be immediately used and trusted by a large number of working neurophysiologists. The ability to process very large datasets (millions of spikes in hundreds of dimensions) in reasonable human and computer time was deemed essential; error rates comparable to those of commonly-used tetrode methods were deemed acceptable. We constructed a suite of three programs enabling tractable analysis of high-count silicon probe recordings, and tested them on data recorded from rat neocortex with 32-site shank electrodes. While traditional spike sorting methods performed extremely poorly on this data, the new algorithms gave close to the theoretically optimal performance. The techniques and software have been developed in a community-led manner, through extensive feedback from a user base of over 200 scientists in 35 neurophysiology labs. It is downloadable and documented at http://klusta-team.github.io/, and is supported by a highly active user-group mailing list.

## Results

The primary challenge of spike sorting with high count silicon probe data arises from the fact that temporally overlapping spikes are extremely common. This phenomenon can clearly be seen by examination of a segment of raw data recorded with high count probes (Figure 1). The spikes seen in this data are diverse, with some detected on only one or two channels, and others spanning large numbers of channels, as expected of pyramidal cells whose apical dendrites are aligned parallel to the shank ^25^. In this data, simultaneous firing of multiple neurons is common. However, simultaneously firing neurons are usually detected on distinct sets of channels. To deal with the problem of temporally overlapping spikes, we therefore pursued an approach which works by extracting spikes as local spatiotemporal events, which are then sorted using a novel cluster analysis algorithm, and manually adjusted using a new computer-guided interface.

**Figure 1.**
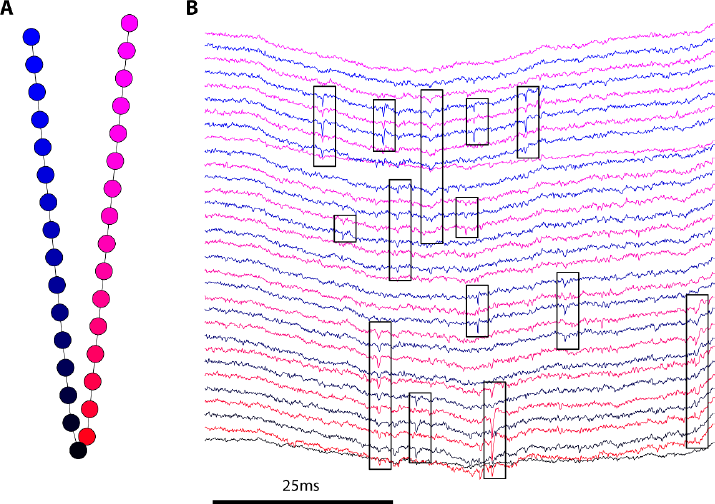
High-count silicon probe recording. **A,** Layout of the 32-site electrode array used to collect test data. **B,** Short segment of data recorded in rat neocortex with this array. Color of traces indicates recording from the corresponding colored site in A. Black rectangles highlight action potential waveforms; note the frequent occurrence of temporally overlapping spikes on separate recording channels.

### Spike Detection

The first step of our spike sorting pipeline is spike detection and feature extraction. This is implemented in the first program of the software suite, *SpikeDetekt*.

SpikeDetekt detects action potentials as spatiotemporally localized events. This step requires knowledge of the probe geometry, which is specified by the user in the form of an “adjacency graph” (Figure 2A). We illustrate the spike detection process with reference to a small segment of data containing two temporally overlapping but spatially separated spikes (Figure 2B).

**Figure 2.**
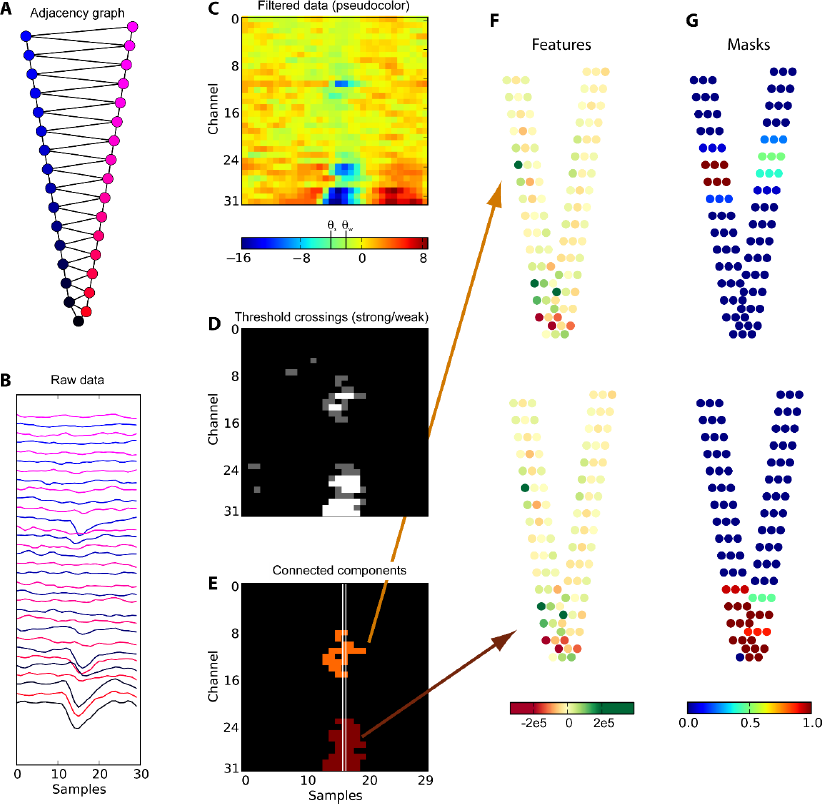
Local spike detection algorithm. **A,** Adjacency graph for the 32-channel probe. **B,** Segment of raw data showing two simultaneous action potentials on spatially separated channels. **C,** High-pass filtered data shown in pseudocolor format. Vertical lines on the colorbar indicate strong and weak thresholds, *θ_s_* and *θ_w_* (respectively 4 and 2 times standard deviation). **D,** Gray-scale representation showing samples which cross the weak threshold (gray), and the strong threshold (white). **E,** Results of two-threshold flood fill algorithm, showing connected components corresponding to the two spikes in orange and brown. Note that isolated weak threshold crossings resulting from noise removed. White lines indicate alignment times of the two spikes. **F,** Pseudocolor representation of feature vectors for the two detected spikes (top and bottom). Each set of three dots represents three principal components computed for the corresponding channel. Note the similarity of the feature vectors for these two simultaneous spikes (top and bottom). **G,** Mask vectors obtained for the two detected spikes (top and bottom). Unlike the feature vectors, the mask vectors for the two spikes differ. Each set of three dots represents the three identical components of the mask vector for the corresponding channel.

The first stage of the algorithm is high-pass filtering the raw data to remove the slow local field potential signal. While a number of approaches could be used, we currently implement a simple linear filter (Butterworth in forward-backward mode). The high-pass filtered signal for the example data segment is shown in pseudocolor form in Figure 2C.

Spike detection works via a double-threshold flood fill algorithm (Figures 2D-E). Two thresholds are defined: a strong threshold, *θ_s_*, and a weak threshold, *θ_w_* (optimal values for these parameters were found to be 4 and 2 times the standard deviation of the filtered signal, as described below). The double threshold flood fill detects spikes as spatiotemporally connected components, in which the filtered signal exceeds the weak threshold for every point, and in which at least one point exceeds the strong threshold. Two points on a single channel are defined to be neighboring if they are separated by one sample in time, and two points at the same time are considered neighboring if they are joined by an edge of the adjacency graph; this allows the algorithm to work not just for linear probes, but probes of any geometry. The dual-threshold approach avoids spurious detection of small noise events, since isolated islands in which only the weak threshold is exceeded are not retained. Conversely, this approach ensures that spikes will not be erroneously split due to noise, as areas joined by weak threshold crossings are merged.

After detection, spikes must be temporally aligned before sorting. Temporal alignment is critical to accurate sorting, as spikes from a single neuron that were poorly aligned could be mistaken for spikes of multiple neurons. We perform spike alignment by assigning each spike a central time given by the center of mass of the spike’s suprathreshold components, weighted by a power parameter *p* (see Methods); note that the central spike time found by this method need not be an integer. Visual inspection showed that spike times detected with this method correspond closely to those that would be assigned by a human operator (Figure 2E). Spike waveforms are realigned to subsample resolution, by resampling around their central time using cubic spline interpolation.

The waveforms of each spike are summarized compactly by two vectors, referred to as the “feature vector” (Figure 2F) and the “mask vector” (Figure 2G). The feature vector is found by applying principal component analysis to the aligned waveforms on each channel (3 principal components were kept for the current analysis). All channels are used in computing the feature vector; thus our two example spikes have very similar feature vectors, as their central times are similar (Figure 2F). The mask vector is computed from the peak spike amplitude on each detected channel, rescaled and clipped so that a channel outside the connected component is given a mask of 0, and a channel with amplitude above *θ_s_* is given a mask of 1. The mask vector allows temporally overlapping spikes to be clustered as separate cells. Indeed, although the feature vectors of our two example spikes were very similar, their mask vectors are completely different (Figure 2G).

### Performance Validation and parameter optimization

To quantify the performance and optimize the parameters of this algorithm requires “ground truth”, augmenting the extracellular recordings with knowledge of when the recorded neurons actually fired. While such ground truth data was obtained for wire tetrode recordings by simultaneous intracellular recording ^8, 9^, no such data currently exists for high-count silicon probes.

We therefore created a simulated ground truth data set, which mimics the most important challenges of the high-count spike sorting problem. To do this, the spike waveform of a “donor cell” identified in one recording, is repeatedly added at known times to a second “acceptor” recording made with same probe. This ensures that the background against which a spike must be isolated is identical to that found *in vivo*; since the extracellular medium is a linear conductor^26^, linear addition of spike waveforms serves as an sufficient model for overlapping spikes. To evaluate the performance of the system, we chose 10 donor cells with a variety of amplitudes and waveform distributions (Figure 3A), again using recordings from rat cortex with a 32-channel probe shank. Finally, in order to model the variability of waveforms produced by a single neuron due to phenomena such as bursting ^27-29^, we scaled each spike to a random amplitude in a range that varied by a factor of 2 (see Methods). We refer to the spikes added to the acceptor data set at known times as “hybrid spikes”, and the resulting data set as a “hybrid data set”.

**Figure 3.**
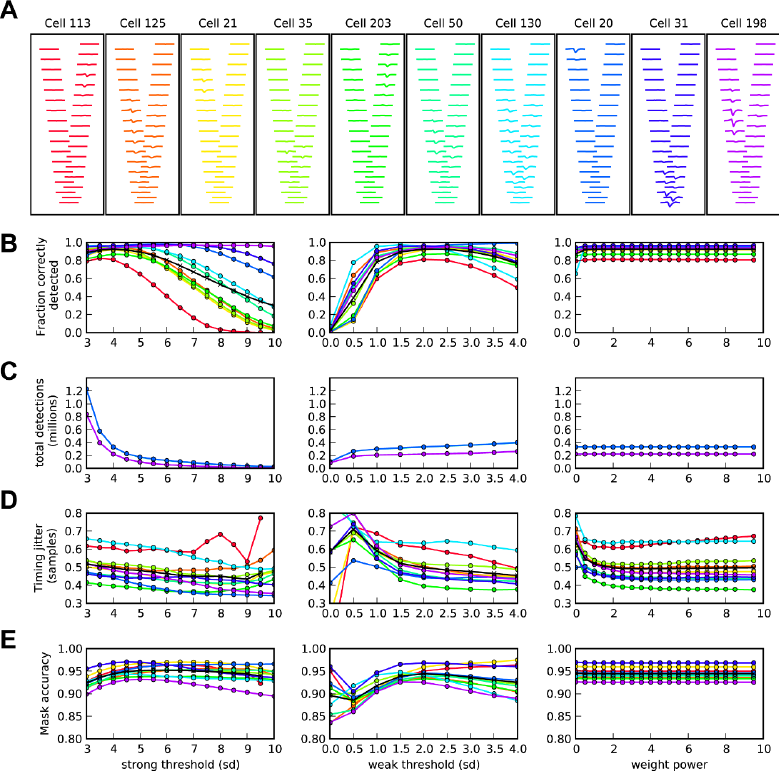
Evaluation of spike detection performance. **A,** Waveforms of the 10 donor cells used to test spike detection performance, in order of increasing peak amplitude (left to right). **B,** Fraction of correctly detected spikes as a function of strong threshold *θ_s_* (left), weak threshold *θ_w_* (center), and power parameter *p* (right). Colored lines indicate performance for the correspondingly colored donor cell waveform shown in A; black line indicates mean over all donor cells. **C, D, E,** Dependence of the total number of detected events, timing jitter, and mask accuracy on the same three parameters.

To evaluate spike detection performance, a heuristic criterion was used to identify which spikes detected by the algorithm corresponded to which hybrid spikes (see Methods). Performance was then measured using four statistics, which were used to evaluate performance as a function of three parameters (*θ_w_*, *θ_s_*, and *p*). The first criterion was the fraction of hybrid spikes detected (Figure 3B). Performance by this measure showed a strong dependence on the thresholds: values of *θ_s_* above 4 times standard deviation (SD) resulted in poor detection, particularly for low-amplitude cells. The dependence of performance on *θ_w_* was more complex: poor performance resulted not just from overly high values (>2.5 SD), but also overly low values (<2 SD). Examination of example errors (not shown) indicated that overly low values of *θ_w_* led to inappropriate merging of temporally overlapping but spatially separated spikes, while overly high values led to artificial splitting of single spikes. Our second measure of performance was the total number of detection events (Figure 3C). Because this includes detections of noise events as well as true spikes of the hybrid and background cells, it is advantageous for this number to be as small as possible provided the fraction correctly detected remains high. We found that this statistic most critically depended on the strong threshold, increasing strongly for values below 4SD.

The third statistic used to evaluate performance was timing jitter, defined as the standard deviation of the difference between the detected and actual times of each hybrid spike (Figure 3D). This statistic showed an improvement for larger values of *θ_s_* and *θ_w_*, reflecting the fact that the most accurate spike times can be estimated from a minority of larger amplitude spikes. Temporal jitter was in all cases less than one sample, and showed consistent variation with the power parameter *p*. For all hybrid cells, jitter was worse for p < 1; for low amplitude cells it showed a further worsening for p>2, reflecting noise introduced by overweighting of peak amplitude times. The final statistic used was mask accuracy, which measures how closely the detected mask vectors match those expected from the ground truth (see Methods). This statistic showed strongest dependence on *θ_w_* with a peak around 2 SD, and less pronounced dependence on *θ_s_* peaking around 5 SD.

We conclude that close to optimal performance can be obtained using a strong threshold of 4 SD, a weak threshold of 2 SD and a power weight of 2. Furthermore, using these parameters yields around 95% correctly detected spikes, and spike timing jitter of 0.5 samples.

## Cluster Analysis

The second step of our spike sorting pipeline is automatic cluster analysis, which is implemented in the program *KlustaKwik*.

For tetrode data, we previously found that cluster analysis using a mixture of Gaussians fit gave close to optimal performance ^8^. This approach cannot be directly ported to high channel count data for two reasons. First, cluster analysis in large dimensions is a notoriously difficult problem due to the “curse of dimensionality”: for any spike, noise measured on the large number of uninformative channels will swamp signals measured on the smaller number of informative channels. Second, because temporally overlapping spikes have similar feature vectors (Fig. 2F), further information such as the mask vectors must be used to distinguish these spikes. While solutions to certain types of high-dimensional clustering problems have been found ^17^, no solution exists to the specific problem that occurs in spike sorting, in which different points are defined by a different subset of relevant dimensions, and millions of points must be clustered in reasonable computer time.

To solve this problem, we designed a novel statistical method for high-dimensional cluster analysis termed the “masked EM algorithm” ^30^. This algorithm fits the data as a mixture of Gaussians, but with each feature vector replaced by a virtual ensemble in which features with masks close to zero are replaced by a noise distribution (see Methods). Channels with low mask values are thus “disenfranchised”, and do not contribute to cluster assignment; furthermore, the probabilistic nature of this disenfranchisement means that false clusters are not created when amplitudes cross an arbitrary threshold. The computational complexity of this algorithm is better than that of the traditional EM algorithm, scaling with the mean number of unmasked channels per spike, rather than the total number of channels. As the number of unmasked channels per spike does not increase for larger electrodes, the algorithm will thus scale to very large electrode arrays.

To evaluate the performance of this algorithm, we used the hybrid data sets described above. For each hybrid data set, we identified the cluster containing most hybrid spikes and computed two numbers: the false discovery rate (fraction of spikes in the cluster that were not hybrids), and the true positive rate (the fraction of all hybrid spikes assigned to the cluster). To estimate the theoretical optimum performance that could be expected, we used the Best Ellipsoid Error Rate (BEER) measure ^8^. The BEER measure fits a quadratic decision boundary using ground truth data, and evaluates its performance with cross-validation; by varying the parameters of the quadric support vector machine used to perform supervised learning, an ROC curve showing optimal performance is obtained. Examination of the masked EM algorithm’s performance on an example hybrid data set showed that performance was close to the theoretical optimum estimated by the BEER measure (Figure 4A). However, the classical EM algorithm’s performance was poor, with error rates typically exceeding 50%. Across all hybrid data sets, we found no significant difference between the total error of the masked EM algorithm and theoretical optimal performance (p = 0.792, t-test), but a significant difference between the performance of the Classical and Masked EM algorithms (p = 0.005, t-test; Figure 4B). To ensure the poor performance of the classical EM algorithm did not simply reflect incorrect parameter choice, we reran it for multiple values of the penalty parameter (which determines the number of clusters found ^30^), but this could not improve Classical EM performance. This analysis also demonstrated that the error rates of the masked EM algorithm were largely independent of the penalty parameter; using a value corresponding to the Bayesian Information Criterion seems a good option for penalty choice, as it led to a reasonably small number of clusters without compromising error rates (Figures 4C, D).

**Figure 4.**
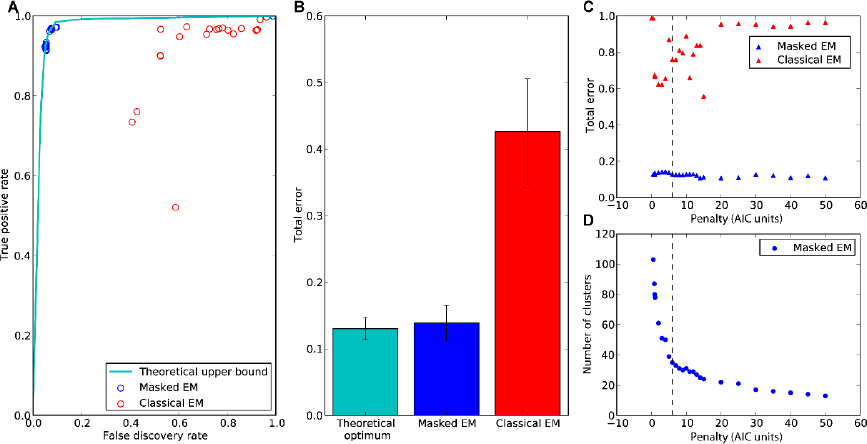
Evaluation of automatic clustering performance. **A.** Receiver-Operating Characteristic (ROC) Curve showing the performance of the Masked EM algorithm (blue) and Classical EM algorithm (red) on one of the 10 hybrid datasets; each dot represents performance for a different value of the penalty parameter. The cyan curve shows a theoretical upper bound for performance, the best ellipsoid error rate (BEER) measure obtained by cross-validated supervised learning. **B,** Mean and standard error of the total error (false discovery plus false positive) over all 10 hybrid datasets for theoretical optimum (BEER measure), Masked EM and Classical EM algorithms. For each data set and measure, the parameter setting leading to best performance was used. **C,** Effect of varying the penalty parameter (as a multiple of the AIC penalty) on the total error for both algorithms. The dotted line indicates the parameter value corresponding to BIC. Note that the Masked EM algorithm performed well for all penalty values. **D,** The number of clusters returned by the Masked EM algorithm as a function of the penalty parameter.

We conclude that the performance of the Masked EM algorithm is close to optimal for this spike sorting problem, yielding false positive and false discovery rates both of the order 5%.

### Manual Validation and Adjustment

The final step of the spike sorting pipeline is manual verification and adjustment of cluster assignments, which are implemented in the program *KlustaViewa*.

Although semi-automatic clustering provides more consistency and lower error rates than fully manual spike sorting ^8^, fully automatic cluster analysis cannot be trusted to accurately classify all spikes, and manual verification is unavoidable. The manual corrections required typically consist of merging clusters corresponding to a single neuron, that were not joined by the automatic algorithm. Erroneous splitting of the spikes of a single neuron most frequently occurs because the waveforms produced by the cell change due to electrode drift, bursting, or other reasons ^27-29^. These waveform shifts are very hard to model and correct mathematically, but can usually be identified by manual inspection of waveforms, auto- and cross-correlograms, and cluster shapes. It is essential that this step be done with a minimum of human operator time, a problem which becomes particularly acute with the very large numbers of neurons recorded by large dense electrode arrays. Specifically, if *N* clusters are produced automatically, it is impractical for a human operator to inspect all *N*^2^ potential merges. We addressed this problem using a semi-automatic guide we termed the “Wizard,” that reduces the number of potential merges that must be evaluated to order *N*.

The Wizard works by presenting the operator with pairs of potentially mergeable clusters, in an order determined by a measure of pairwise cluster similarity. Because the Wizard is used in an iterative fashion, it is essential that this similarity measure be computable in a fraction of a second, even for datasets containing millions of spikes. Thus, only metrics based on the summary statistics of each cluster, rather than individual points, are suitable. We evaluated several candidate similarity measures. The Kullback-Leibler divergence between two Gaussian distributions was not a suitable as it overweighted differences in covariance matrix relative to differences in the mean. However, we found that good performance was obtained by using a single step of the EM algorithm to compute the similarity of the mean of one cluster to each of the others (Figure 5A). To verify the accuracy of this measure, we simulated automatic clustering errors by splitting the ground truth clusters in the hybrid data sets into two subclusters containing high and low amplitude spikes. In all cases, the similarity measure correctly identified the other half of the artificially split cluster (Figure 5B).

**Figure 5.**
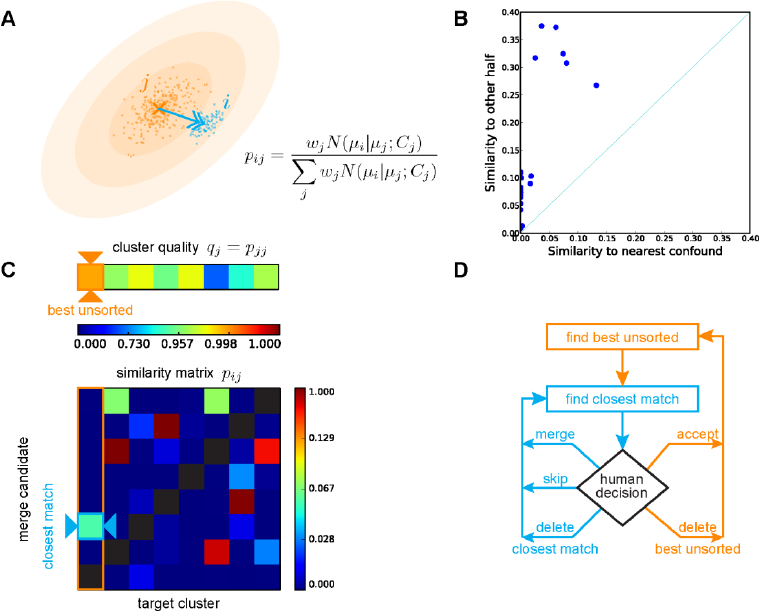
The “Wizard” for computer-guided manual correction. **A,** Illustration of the measure used to quantify cluster similarity. *p_ij_* represents the posterior probability with which the EM algorithm would assign of the mean of cluster *i* to cluster *j*. **B,** To test this measure, the clusters corresponding to hybrid spikes were artificially cut into halves of high and low amplitude. In each case, the similarity measure identified the second half as the closest merge candidate. **C,** The wizard identifies the best unsorted cluster as the one with highest quality (top), and finds the closest match to it using the similarity matrix. **D,** The wizard algorithm. The best unsorted cluster and closest match are identified. The operator can choose merge the closest match into the best unsorted, ignore the closest match, or delete it by marking it as multiunit activity or noise; the wizard then presents the next closest match to the operator (blue arrows). After a sufficient number of matches have been presented, the operator can decide that no further potential matches could have come from the same neuron. The operator can then accept the best unsorted cluster as a well-isolated neuron, or delete it as multiunit activity or noise. The wizard then finds the next best unsorted cluster to present to the operator (orange arrows).

The Wizard was implemented in a new graphical software tool designed specifically for manual-stage sorting of high-count probe data, that we term KlustaViewa. The manual stage can take several hours of operator time, and operator errors are lowest in the start of this period. The Wizard was therefore designed to present the operator with decisions that can be made quickly, with the most important decisions presented first. It does this using a two-level iterative method (Figure 5C). The Wizard iterates through all clusters starting with the best currently unsorted spikes. The remaining clusters are ordered by similarity to the best unsorted cluster, and the decision of whether to merge, split, or delete each merge candidate is in turn made by the operator (Figure 5D). Once satisfied that no more potential merges exist for the currently best unsorted cluster, for the operator decides to accept it as a well-isolated neuron or reject it as multiunit activity or noise, and the top-level iteration begins again. Note that while the Wizard guides the operator through the decision process, the operator at all times has easy access to all data required to make a rapid decision on merging, provided by KlustaViewa’s user-friendly and easily-navigable graphical user interface (Figure 6). This process allows for thorough manual verification of a dense-array recording in just a few hours.

**Figure 6.**
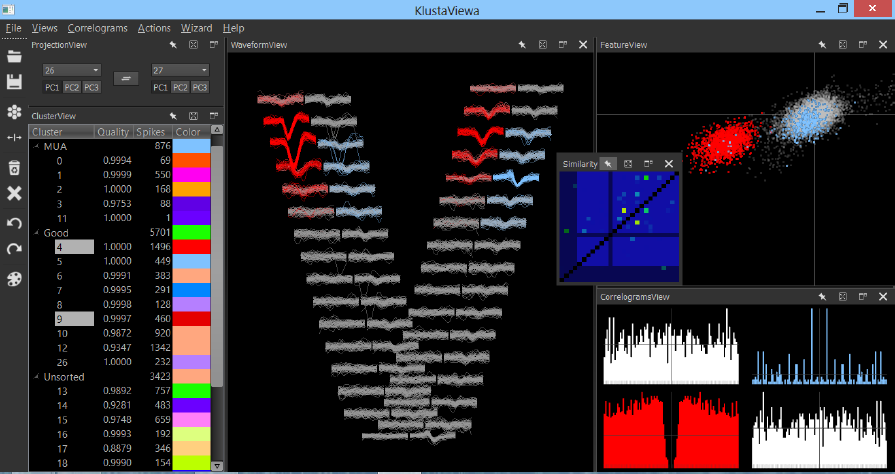
Screenshot of the KlustaViewa graphical user interface.

## Discussion

We have produced a software suite for spike sorting of data from large, dense electrode arrays. Analysis of simulated ground-truth data indicated that error rates of this approach are of the order 5%.

A critical step in this system, and all other spike sorting systems currently in wide use, is manual correction and verification. While a system that could detect and classify action potentials fully automatically would be an extremely useful tool, the complexity of the physical processes underlying spike waveform variability makes this “artificial intelligence problem” currently intractable. Extracellular array recordings are subject to numerous sources of error including electrode drift, overlapping spikes, and the fact that neuronal spike waveforms are not constant, but change according to firing patterns including but not limited to bursting ^27-29^. While most working neurophysiologists have a good understanding of these potential artifacts, formalizing this knowledge into a reliable mathematical model has proved challenging. Because spike sorting errors could lead to erroneous scientific conclusions ^29^, it remains essential that a scientist is able to inspect the results produced by an automatic algorithm, then correct or discard its results. This situation is common in many fields of high-throughput biology, for example connectomics ^14^.

The current performance of the system is sufficient for practical analysis of large-scale electrophysiological recordings. Nevertheless, there remain areas for further improvement. The first of these concerns execution time, for which the most critical component is the clustering algorithm. The KlustaKwik program is highly optimized, and several orders of magnitude faster than standard mixture of Gaussians fitting. Nevertheless, when running on very large data sets (millions of points in hundreds of dimensions), it can take hours or even days of computer time to complete. We are currently investigating two approaches to speed up this analysis stage: first, by hardware acceleration such as GPU ^31^ or cloud computing ^32^; and second, by examining whether cluster analysis algorithms that exclude the most computationally expensive step of covariance matrix estimation, can give adequate performance (e.g. Refs.^33, 34^). A second area for improvement regards the detection of spatiotemporally overlapping spikes. While the majority of temporally overlapping spikes occur on distinct sets of channels, spikes that overlap in both space and time cannot be resolved by the current algorithm. Template-matching algorithms have solved this problem in the case of *in vitro* retinal array data ^35, 36^ but this data is much less noisy than *in vivo* brain recordings. While recent research suggests that certain forms of template matching may succeed at least for tetrode data *in vivo* ^18, 21^, this approach has yet to gain the confidence of the experimental community, and numerous challenges of execution speed and manual verification need to be overcome. The platform we have described here constitutes a starting point for the development of practical *in vivo* spike sorting methods based on template matching.

Our evaluation of spike sorting performance was based on synthetic ground truth data as no simultaneous recordings of large extracellular arrays with unambiguous single neuron identification yet exist. The additional spike variability that can occur for example due to bursting was modeled by a random amplitude scaling. Nevertheless, biological neurons may show more complex waveform variability, as well as artifacts such as electrode drift that so far must be corrected manually. Such data may become available in the future, for example by combination of large-scale silicon probe recording data and temporally improved optical methods. The resulting data is likely to lead to further improvements in spike sorting performance.

Research in neural probe technology is advancing rapidly, and will soon allow production of probes allowing simultaneous recording from hundreds of closely-spaced recording sites. Our system was tested using data from a 32-site shank, the largest currently available to us. However, because our algorithms are based on local analysis, their computational complexity scales with to the number of unmasked features per spike, rather than the total number of recorded channels. Thus, the performance and execution speed reported here is also expected for the forthcoming generation of very large neural electrode arrays.

## Methods

### Test data

To test the algorithm, we created simulated ground truth data using a method termed “hybrid data sets”. The raw data used to construct this ground truth consisted of two separate recordings from somatosensory cortex (−3.8 mm from bregma, 3 mm lateral to midline, 1mm depth) of sleeping adult rats, using 32-site silicon probes in a staggered configuration mounted on a home-made microdrive. Ground and reference electrodes were stainless steel screws over the cerebellum. Data was continuously recorded wideband (1Hz-Nyquist), at a sampling rate of 20 kHz.

To create the hybrid data sets, we first completed a full spike sorting of each data set, including manual verification. Five clusters were chosen from each data set, corresponding to neurons spanning the range of amplitudes and channel distributions observed in the data (Figure 3A). The mean unfiltered waveform of each neuron was computed, its mean was subtracted, and its value at each end was set to exactly zero by tapering with a hamming function. These “donor waveforms” were added at prescribed times to the raw unfiltered data of the other “acceptor” recording. To simulate amplitude variability, we linearly scaled each added waveform by a random factor chosen from the range 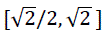 causing amplitudes to vary by a factor of two, which suffices to capture the variability typical of bursting neurons ^27^. The interspike intervals typical of bursting neurons were not simulated as this does not affect the spike detection or clustering process; instead, hybrid spikes were added regularly at rates in the range 2–4 spikes per second. To ensure that the simulated data tested the ability of our software to realign spikes to subsample resolution, each added spike was shifted by a random subsample offset using cubic spline interpolation.

### File format

To implement the software, we designed an HDF5-based file format to store raw data, intermediate analysis results (such as extracted spike waveforms and feature vectors), as well as final data such as spike times and cluster assignments ^37^. The format makes use of HDF5 links to allow a single, small file (the “.kwik file”) containing all data required for scientific analysis (e.g. spike times, cluster assignments, unit isolation quality measures). Bulky raw data and intermediate processing steps such as feature vectors are stored in separate files (the “.kwd” and “.kwx” files). This “detachable” format is designed for data sharing applications, allowing users to download as much data as required for their needs. A full specification of the format can be found at https://github.com/klusta-team/kwiklib/wiki/Kwik-format.

### SpikeDetekt

Spike detection was implemented by SpikeDetekt, a custom program written in Python 2.7 using the packages NumPy, SciPy, and PyTables.

The first step of the program is to filter the raw voltage trace data to remove the low-frequency local field potential (LFP). This is achieved with a 3rd order Butterworth filter used in the forward-backward mode to ensure zero phase distortion. Filter parameters can be specified by the user; for the analyses described here we used a band-pass filter of 500 Hz to 0.95*Nyquist.

The second step is threshold determination. Spike detection thresholds are specified as multiples of the standard deviation of the filtered signal; at the option of the user, a single threshold is used for all channels in order to avoid emphasizing noise from low-amplitude channels. To boost execution speed while minimizing the chance of biased estimates, the standard deviation is estimated from five data chunks of length 1 second each, picked randomly from throughout the recording. The standard deviation is computed with a robust estimator, median(|*V*|)/.6745, to avoid contamination by spike waveforms.

The next step is spike detection. The spike detection code operates on consecutive chunks of data (1s length) for memory efficiency. Spatiotemporally connected regions of weak threshold crossing are detected using a non-recursive flood fill algorithm, with spatial continuity defined using a user-specified adjacency graph. Only connected components for which at least one point exceeds the strong threshold are kept for further analysis.

Spike alignment is computed based on a scaled and clipped transformation of the filtered voltage *V*(*t*, *c*):

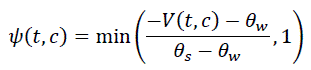

Note that *ψ*(*t*, *c*) can never be negative within a spike, as the floodfill algorithm only finds points for which −*V*(*t*, *c*) > *θ_w_*. The center time for each spike *S* is computed as

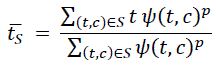

where (*t*, *c*) ∈ *S* denotes the set of times and channels, for all points assigned to this spike by the floodfill algorithm. If *p* = 1, this formula measures the spike’s center of mass; if p = ∞, it measures the time of the spike peak.

Spikes were realigned on *t̄_S_* to subsample resolution using cubic spline interpolation (note that the center time will in general not be an integer number of samples). Feature vectors are computed for each channel separately by principal component analysis; the number of features per channel is a user settable parameter, with default value 3. Finally, mask vectors are computed for each spike *S* as zero for channels not appearing in the connected component, and as the maximum scaled waveform for all channels inside the component:

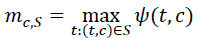

To evaluate the performance of SpikeDetekt, required identifying which detected spikes correspond to ground truth spikes. This was done with a dual criterion: the difference between the detected time and ground truth needed to be less than 2 samples, and the detected mask vector **m_s_** needed to have a similarity to the ground truth mask vector **m_G_** of at least 0.8, defined by the mask similarity measure

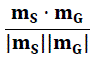

Note that mask similarity cannot exceed 1, by the Cauchy-Schwartz inequality. The validity of this criterion was verified by showing that detected spike timing jitter rapidly increased for similarity threshold for values less than 0.8, but was insensitive to threshold value above 0.8. Once the detected spikes corresponding to ground truth had been identified, the four measures in figure 3 were computed. This analysis used the Python library Joblib to prevent unnecessary recomputation.

### KlustaKwik

Automatic clustering was performed by KlustaKwik, a custom program written in C++. The first version of this program was designed for tetrode data, implemented a hard EM algorithm for maximum-likelihood fitting of a mixture of arbitrary-covariance Gaussians, and was released in 2000 but not specifically described in a published manuscript. Here, we have implemented several modifications of this software to enable automatic sorting of high-count probe data. The program now implements a novel “masked EM algorithm” ^30^ designed for high dimensional classification, as well as other features such as cache optimization resulting in a speed increase of over 10,000%.

The masked EM algorithm takes as input both feature vectors and mask vectors. It works by fitting a mixture of Gaussians to a virtual data set in which each feature vector is replaced by a probability distribution:

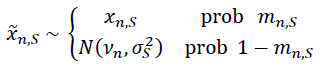

Here, *x_n, S_* represents the *n*^th^ component of the feature vector for spike *S*; *m_n,S_* represents the *n*^th^ component of the mask vector for spike *S*; and 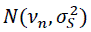 denotes a univariate Gaussian distribution with mean and variance equal to those of the subthreshold noise distribution of the *n*^th^ feature.

The masked EM algorithm consists of alternation of an “E step” in which each spike is assigned to the cluster for which it has highest posterior probability, and an “M step” in which the means and covariances of each cluster are estimated. We have derived analytic formulas for the expectation of the cluster assignment probability used in the E-step, and the cluster mean and variance used in the M step, over the virtual probability distribution *x̃_n,i_* ^30^. Thus, explicit sampling from the virtual distribution does not need to be performed; furthermore, these expectations can be computed much faster than those of the full EM algorithm as they scale with the square of the number of unmasked features, rather than the square of the total number of features.

KlustaKwik automatically determines the number of clusters that best fit the data, determined using a penalty function that encodes a preference for fits with smaller numbers of clusters. We have found a modification of the Bayesian Information Criterion to deal with masked data works well in practice ^30^. Because the algorithm allows for dynamic splitting and merging of clusters during the fitting process, a search for the optimal number of clusters can be achieved in a single run of the algorithm. We have found that starting the algorithm from an initial clustering determined heuristically from the mask vectors avoids the problem of local maxima, and allows good results to be obtained from a single run.

### KlustaViewa

Manual correction of automatic clustering is performed with KlustaViewa, a custom program written in Python 2.7. The manual stage requires interactive visualization of very large numbers of data points, for which existing libraries such as matplotlib were not suitable. We therefore designed a new Python library for rapid interactive data visualization named Galry ^38^. Galry leverages the computational power of modern graphics processing units ^31^ through the OpenGL graphics library ^39^. High performance is achieved by porting most visualization computations to the GPU using custom shaders, and by minimizing the number of OpenGL API calls through batch rendering techniques.

To ensure rapid adoption by the experimental community, we designed KlustaViewa’s user interface by the integrating novel features necessary for high-count probes into a user interface as similar as possible to existing manual spike sorting environments such as Klusters ^13^. In addition to data views familiar from previous spike sorting systems (such as waveform, auto- and cross-correlograms, and similarity matrix), we implemented several new features. The most important of these is the Wizard (described in the main text), that automatically leads the user through the manual verification and merging process, while always allowing the user free access to all of the views familiar from standard spike sorting systems. In addition, a number of enhancements were designed specifically to make the sorting of high-count probe data tractable. These include features to allow display of masking information; rapid and automatic display of the channels relevant to selected clusters; transient color brushing ^40^; and automatic downsampling to ensure low latency display when dealing with very large data sets.

The Wizard is based on a metric of similarity for each pair of clusters. This was computed by running a single step from the EM algorithm to compute the posterior probability for assigning the mean of cluster *i* to cluster *j*:

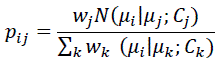

Here *w_j_* represents the weight of cluster *j* (i.e. the fraction of points already assigned to this cluster); *µ_j_* and *C_j_* represent its mean and covariance as computed by the M-step of the masked EM algorithm. The quality of each cluster *j* was defined as the diagonal element *p_jj_*, i.e. the posterior probability for classifying cluster *j*’s mean as coming from cluster *j* itself. A high value for *p_jj_* therefore indicates that cluster *j* has no close neighbors.

## Contributions

C.R., D.F.M.G., S.N.K. and J.S. wrote SpikeDetekt. K.D.H, S.N.K., and D.F.M.G. designed the Masked EM algorithm and wrote KlustaKwik. C.R. wrote KlustaViewa and Galry. S.N.K. analyzed algorithm performance. M.A.B. and G.B. recorded the test data. K.D.H., S.N.K., and C.R. wrote the manuscript with inputs from all authors.

## Acknowledgements

We thank the 100+ members of the mailing list for their feedback, bug reports, and suggestions. This work was supported by EPSRC (K015141, I005102) and the Wellcome Trust (95668).

